# In vivo modulation of network activity drives the nanoscale reorganisation of axo-axonic synapses at the axon initial segment

**DOI:** 10.1101/2024.11.29.625981

**Authors:** B. Compans, V. Mastrolia, C. Lenherr, J. Burrone

**Author notes:** These authors contributed equally to this work (B. Compans, V. Mastrolia).

## Abstract

Chemical synapses control their strength through the nanoscale clustering of postsynaptic receptors into sub-synaptic domains (SSDs). Despite their importance in synapse function, the properties and plasticity of these domains are not well understood *in vivo*, particularly in inhibitory synapses. We used direct Stochastic Optical Resolution Microscopy (dSTORM) to show that Gephyrin, the main inhibitory receptor scaffold protein, is organised into SSDs *in vivo*, with distinct arrangements depending on their sub-cellular location and presynaptic partner. Furthermore, chronic chemogenetic increases in cortical activity caused a reduction in Gephyrin SSD volume specifically in axo-axonic, but not axo-dendritic, synapses. Functionally, this resulted in a weakening of axo-axonic contacts. We show that the nanoscale arrangement of synapses in the brain is plastic and used to fine-tune synaptic gain *in vivo*.

## Introduction

The ability to image molecules with nanoscale precision has uncovered novel arrangements within cells that have revolutionised our understanding of how structure determines function (1). This is particularly the case at the chemical synapse, where transmission of information is thought to be defined by the local, nanoscale distribution of synaptic proteins. Recent work has shown that synapses have very precise sub-synaptic arrangements that can only be resolved with super-resolution microscopy, and which are thought to be important for their function. At excitatory synapses, for example, both the scaffolding proteins at the postsynaptic density and the AMPA receptors that sense glutamate form tight sub-synaptic domains (SSDs) that align with presynaptic clusters of active zone proteins to form a nanocolumn that spans the synaptic cleft (2–5). Similarly, inhibitory synapses show that the scaffolding protein Gephyrin and the GABA receptors that they tether, are also arranged in SSDs (6–12). It is thought that the overall strength of a synapse can be gleaned from the postsynaptic arrangement - density and area - of receptors within a sub-synaptic domain (2, 13–15). However, although suggested in glutamatergic synapses (4, 16–18), whether subsynaptic domains in GABAergic synapses are present in intact tissue and whether they play a role in modulating synaptic gain *in vivo* remains unknown.

Pyramidal neurons in the cortex receive inhibitory inputs from a heterogeneous population of GABAergic interneurons that target distinct sub-cellular domains and therefore modulate pyramidal cell function in different ways (19). Whereas dendritic GABAergic inputs, typically formed by somatostatin-expressing interneurons, locally influence the integration of excitatory inputs along dendritic segments, interneurons targeting the axon-initial segment (AIS), known as chandelier cells, are thought to directly modulate neuronal output. Although the precise subcellular location of a GABAergic synapse defines the type of modulation it imparts on pyramidal cells, whether the nanoscale arrangement of GABAergic synapses varies along different sub-cellular compartments is not known.

Both excitatory and inhibitory synapses undergo activity-dependent forms of plasticity that modulate synaptic strength. Recent work in excitatory synapses has shown that synaptic plasticity also takes place at the nanoscale (4, 5, 20–25). For example, NMDA-dependent increase in spine size correlates with an increase in SSD number, suggesting SSDs act as modular units that are added or removed by experience-dependent plasticity (5). Together with the dynamics of glutamate receptors at the surface, these nanoscale molecular arrangements are thought to control the strength of synaptic connections. However, whether synapses are plastic at the nanoscale level *in vivo* remains to be determined. Here, we tackle this question by focusing on GABAergic synapses that form onto two distinct subcellular compartments of L2/3 pyramidal neurons in the somatosensory cortex: axo-axonic synapses at the AIS, and axo-dendritic synapses on dendritic tufts. We found that GABAergic synapses show specific SSD arrangements, depending on their subcellular location and the identity of their presynaptic partner. Furthermore, we used a chemogenetic approach to increase network-wide activity in the mouse somatosensory cortex over 2 days (from P20 to P22) to assess activity-dependent forms of plasticity at the nanoscale. We found that increases in activity had no effect on the number of axo-axonic synapses along the AIS but did cause a strong reduction in Gephyrin SSD volume, resulting in a functional weakening in the connections between chandelier cells and pyramidal neurons. Axo-dendritic synapses, on the other hand, remained unaltered, suggesting a further layer of specificity in the rules that drive the plasticity of different inhibitory synapses. Our findings show that the nanoscale plasticity of GABAergic synapses does take place *in vivo* and is responsible for modulating synaptic gain.

## Results

We labelled GABAergic postsynaptic compartments by using intrabodies that specifically recognise Gephyrin (Fibronectin intrabodies generated with mRNA display - FingR) (26), the main scaffolding protein that tethers GABA receptors to the synapses. When fused to EGFP (Gephyrin-FingR-EGFP) and expressed together with cytoplasmic tandem-dimer tomato (tdT) in layer 2/3 pyramidal neurons of the somatosensory cortex, we were able to clearly identify puncta all along the AIS (Fig 1A-B). These puncta co-localised with presynaptic terminals labelled with vGAT (Fig 2D-E, Supp Fig. 1A). We saw clear labelling of synapses along the entire dendritic arbour, although these puncta were typically found at much lower density in dendrites (27).

**Fig 1.**
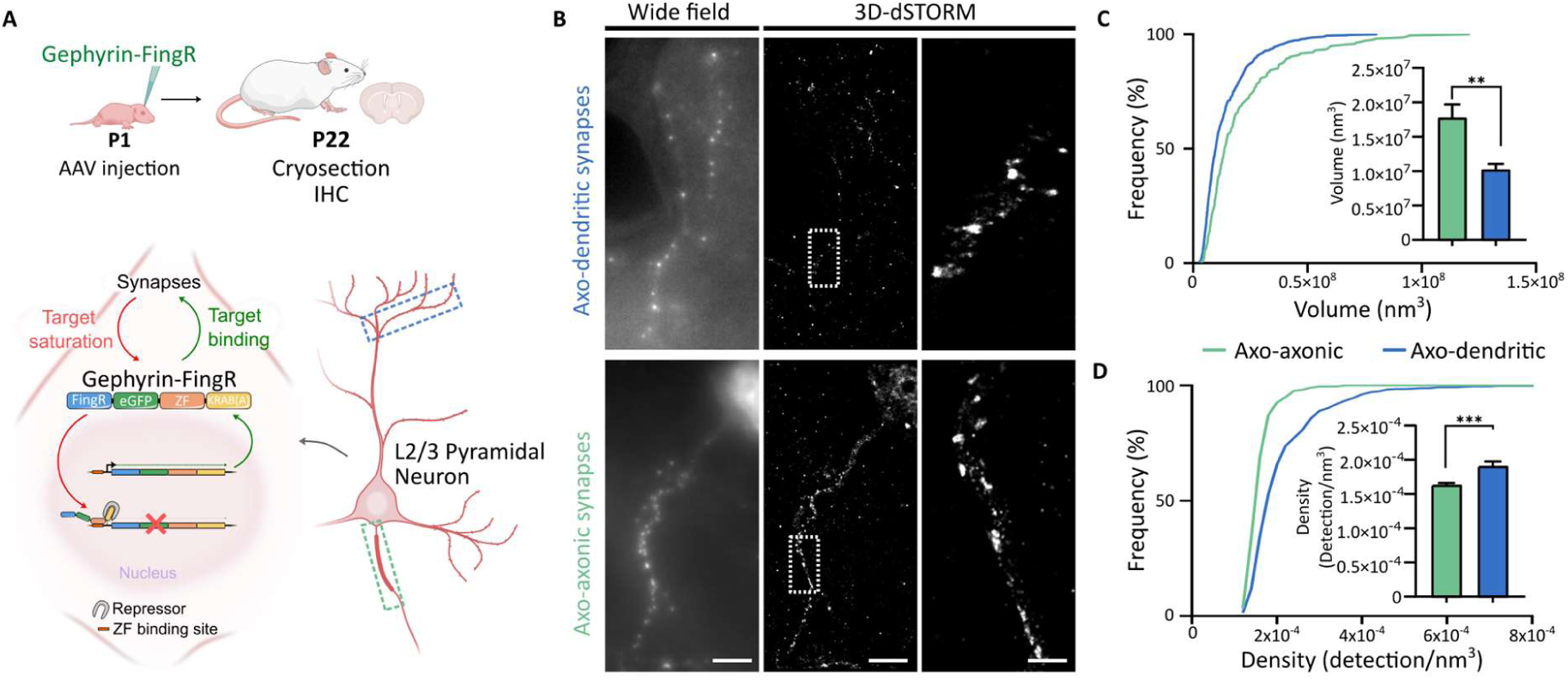
The nanoscale arrangement of inhibitory synapses shows input specificity. (A) Expression strategy for the gephyrin-targeting FingR. (B) Widefield and 3D-dSTORM images of gephyrin-FingR-labelled axo-axonic and axo-dendritic synapses of L2/3 pyramidal neurons imaged in either L2/3 and L1, respectively. (C) Cumulative distribution of gephyrin SSD volume between axo-axonic and axo-dendritic synapses. Medians per cell are shown in the inset as mean +/- SEM, statistics with non-parametric t-test with Mann-Whitney (axo-dendritic n = 21, axo-axonic n = 17, p < 0.0001). (D) Cumulative distribution of gephyrin densities per SSD between axo-axonic and axo-dendritic synapses. Medians per cell are shown in the inset as +/- SEM, statistics with non-parametric t-test with Mann-Whitney (axo-dendritic n = 21, axo-axonic n = 17, p < 0.0001).

**Fig 2.**
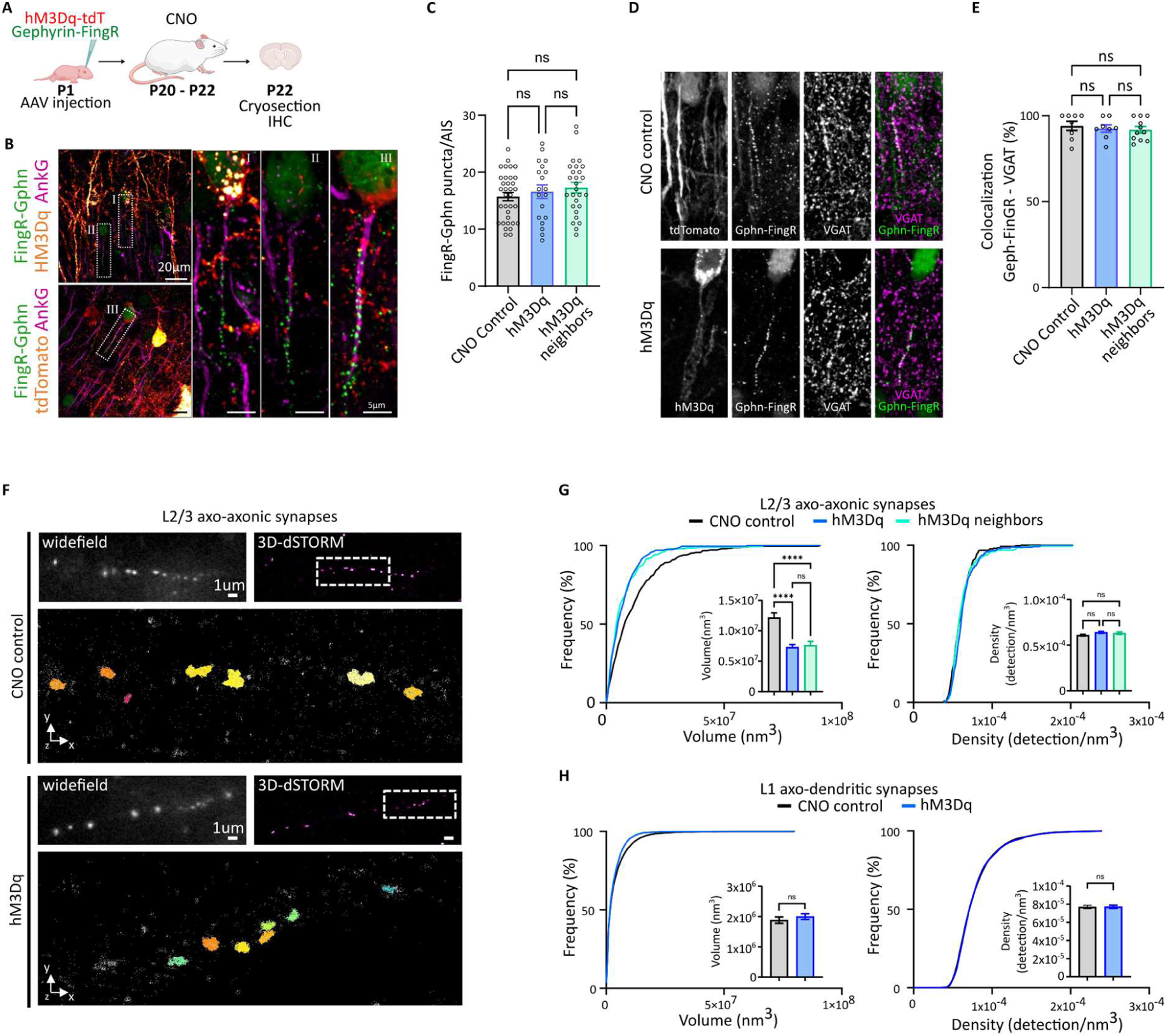
Elevated network activity causes an input-specific decrease in SSD volume of axo-axonic synapses. (A) Expression strategy for the gephyrin-targeting FingR and of the activating DREADD hM3Dq. (B) Confocal microscopy images of axo-axonic synapses (gephyrin-FingR, green) and AISs (AnkG, magenta) in hM3Dq-mCherry and tdT (red) expressing pyramidal neurons. Panels BI-II and panel BIII show example AISs from hM3Dq-positive or tdT-positive neurons, respectively. (C) Quantification of gephyrin-FingR puncta per AIS lengths from neurons expressing tdTomato+ (CNO control), hM3Dq+ or hM3Dq- neighbouring neurons. Means are shown +/- SEM, statistics with One-way ANOVA test (CNO controls n = 8, hM3Dq+ n = 8, hM3Dq- neighbours n = 11, p < 0.0001). (D) Confocal microscopy images from layer 2/3 of S1 cortex showing axo-axonic synapses labelled with gephyrin-FingR and GABAergic presynaptic terminals labelled with vGAT, in hM3Dq or tdTomato-expressing pyramidal neurons. (E) Quantification of gephyrin-FingR and vGAT colocalization from neurons expressing tdTomato+ (CNO control), hM3Dq+ or hM3Dq- neighbouring neurons. Means are shown +/- SEM, statistics with One-way ANOVA test (CNO controls n = 8, hM3Dq+ n = 8, hM3Dq- neighbours n = 11, p < 0.0001). (F) Widefield and 3D-dSTORM images of gephyrin-FingR SSDs in axo-axonic synapses from L2/3 pyramidal neurons of S1 cortex expressing either hM3Dq-tdT or tdT alone, both treated with CNO. (G) Cumulative distribution and means +/- SEM (inset) of gephyrin-FingR SSD volume (left) and density per SSD (right), from axo-axonic synapses belonging to L2/3 pyramidal neurons expressing tdTomato+ (CNO controls), hM3Dq+ or hM3Dq- neighbouring neurons. Means are shown +/- SEM, statistics with One-way ANOVA test (CNO controls n = 21, hM3Dq+ n = 17, hM3Dq- neighbours n = 18, p < 0.0001). (H) Cumulative distribution and means +/- SEM (inset) of gephyrin-FingR SSDs volume (left) and density per SSD (right), from axo-dendritic synapses in L1 belonging to pyramidal neurons expressing tdTomato+ (CNO controls) or hM3Dq+. Means are shown +/- SEM, statistics with non-parametric t-test with Mann-Whitney (CNO controls n = 16, hM3Dq+ n = 17, p < 0.0001).

To explore the nanoscale organisation of these postsynaptic compartments, we carried out direct Stochastic Optical Resolution Microscopy (dSTORM) of Gephyrin-FingR-EGFP puncta in thin slices of fixed somatosensory cortex tissue (Fig 1B-D). We found that we were able to reliably reconstruct the arrangement of gephyrin molecules in both axo-dendritic, typically innervated by somatostatin interneurons (Fig 1B-top), and axo-axonic synapses, typically innervated by ChCs (Fig 1B-bottom), with high spatial resolution. Whereas axo-axonic synapses were imaged at the AIS, axo-dendritic synapses were imaged in distal dendrites, specifically in the tuft region found in layer 1. We found that in both locations, gephyrin puncta were clustered in SSDs that varied in volume and packing density (Fig 1C-D). Importantly, in hippocampal cell cultures, we show that these SSDs correlated well with the clustering of GABA_A_ receptors, suggesting gephyrin SSDs represent a reliable measure for the arrangement of inhibitory synapses and their function (Supp Fig.4). At axo-axonic synapses, Gephyrin SSDs had a larger volume and a lower density of molecules than axo-dendritic synapses (Fig 1C-D), pointing to a previously unknown level of specificity in the nanoscale organisation of inhibitory synapses along different sub-cellular compartments. In line with this, we also observed axo-axonic synapses had larger presynaptic boutons, when labelled with vGAT, than axo-dendritic boutons (Supp Fig. 1B). Within the AIS, Gephyrin SSDs were highly heterogeneous and both the density and volume of Gephyrin SSDs varied in a systematic manner with distance from the soma (Supp Fig. 2C-D), suggesting interesting levels of sub-AIS organisation. We conclude that GABAergic synapses in intact tissue have subsynaptic domains with distinct properties depending on their location within a cell and the identity of the presynaptic interneuron that innervates them.

To assess whether activity-dependent forms of plasticity modulate inhibitory synapses, particularly at the nanoscale, we used a chemogenetic approach to increase network activity in the somatosensory cortex at specific times during development (Fig 2A). Mice were co-injected at P1 in L2/3 of the somatosensory cortex with AAVs expressing Gephyrin-FingR-EGFP and hM3Dq-mCherry, the chemogenetic receptor used to activate pyramidal neurons. We restricted the expression of hM3Dq-mCherry to pyramidal neurons by using a CaMKII promoter. To induce the hM3Dq-mediated activation of neurons, we treated mice with Clozapine N-oxide (CNO) twice daily from P20 to P22, a time window that lies immediately after the period of synapse formation (Fig 2A). Control animals were injected with AAVs for the expression of tdT, instead of hM3Dq, and still treated with CNO. We assessed the levels of network activity by staining for the immediate early gene cFos (Supp Fig. 3A). As described in previous work (28), CNO resulted in a strong increase in activity across the entire network (Supp Fig. 3B). However, this robust increase in network activity from P20 to P22 did not result in a change in axo-axonic synapse number (Fig 2B-C), in contrast to the large reduction in axo-axonic synapse number observed when activity was increased just a few days earlier, during the period of synaptogenesis (P12-P18) (28). We did, however, observe a shortening in the length of the AIS, not just in cells expressing hM3Dq, but also in their hM3Dq-negative neighbouring cells (Supp Fig. 3D). This type of activity-dependent plasticity has been previously described *in vivo* in response to changes in network activity (29,30).

Finally, we investigated whether these nanoscale forms of plasticity had any functional consequence on the strength of axo-axonic connections (Fig 3). We first artificially interfered with Gephyrin clustering to assess its consequences on GABA_A_ receptors. Increasing or decreasing Gephyrin SSD volume in hippocampal neuronal cultures, through either the overexpression of wild type Gephyrin or a dominant-negative form of Gephyrin that interferes with Gephyrin clustering (31), respectively, also resulted in a corresponding change in GABA_A_ receptor SSD volume (Supp Fig. 5A-B). This effect was seen in both axo-dendritic (Supp Fig. 5A-B) and axo-axonic synapses (Supp Fig. 5C-D), suggesting manipulations that alter Gephyrin volume also alter GABA_A_ receptor clustering. To directly measure GABA_A_ receptor function at axo-axonic synapses following *in vivo* network hyperactivity, we carried out targeted patch-clamp recordings from pyramidal neurons in acute slices obtained from a mouse line in which ChCs expressed channelrhodopsin-2 (ChR2) (Fig 3A). Optogenetic activation of ChCs (5 light pulses delivered at 20 Hz) elicited evoked postsynaptic currents (IPSCs) that showed weak depression between consecutive pulses (Fig 3B-D). More importantly, when patching pyramidal neurons in a network that had undergone hyperactivity, by expressing hM3Dq-mCherry and administering CNO in the P20-P22 time window, we found that the amplitude of IPSCs was significantly smaller than the mCherry-only controls (Fig 3C). We saw some short-term plasticity changes (Fig 3D), but no changes in failure rate (Supp Fig. 6K), suggesting a mostly postsynaptic modulation. Since we observed no change in synapse number (see Fig. 2B-E), we concluded that the nanoscale reduction in subsynaptic domain volume was responsible for the decrease in IPSC amplitude. In addition, having observed a decrease in the length of the AIS in hyperactive networks, we also measured the firing properties of pyramidal neurons. However, we saw no major differences in intrinsic excitability, apart from a lower firing frequency at the maximal current injection in hM3Dq-positive neurons (Supp Fig. 6A-D).

**Fig 3.**
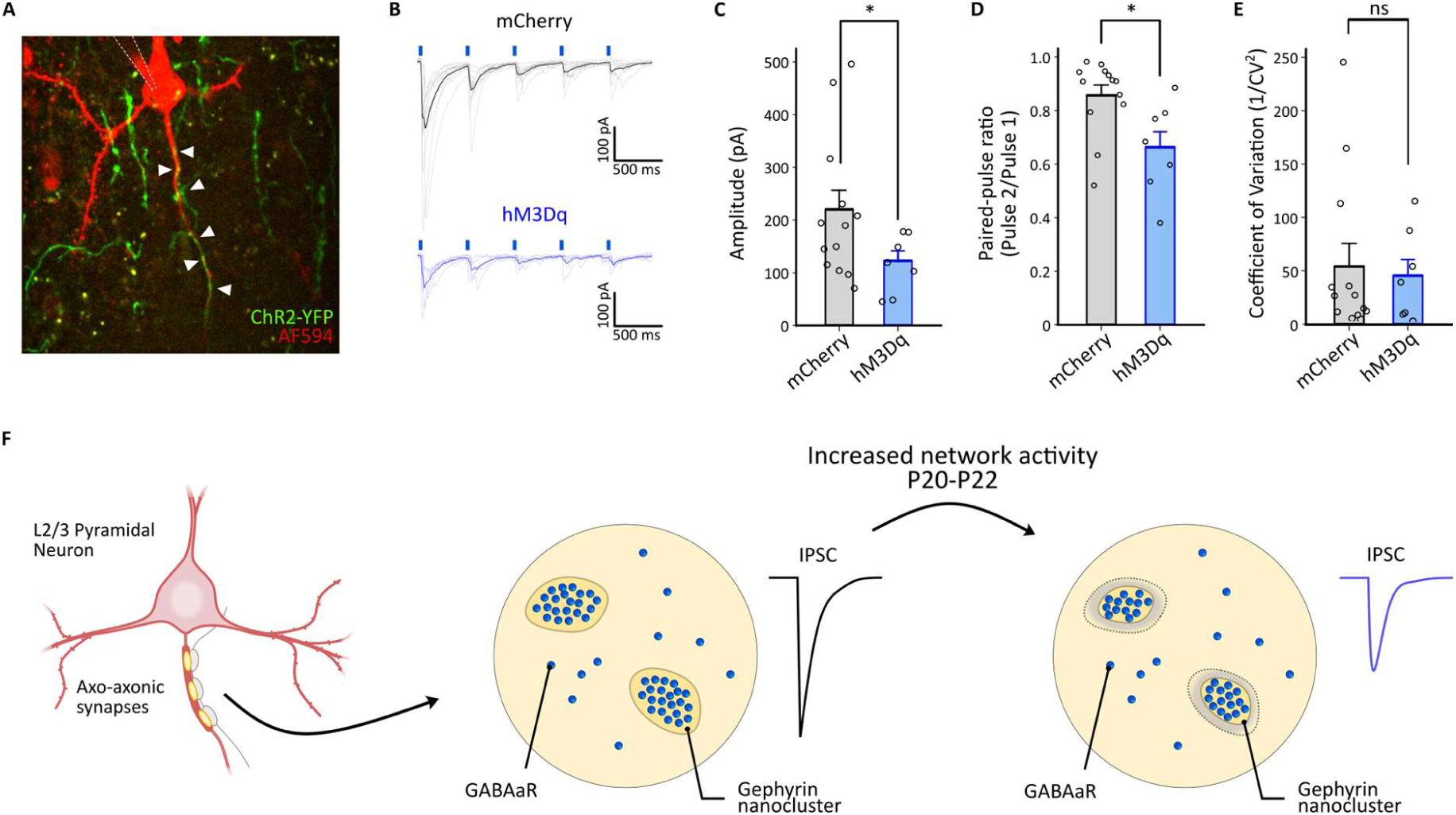
Increased network activity weakens axo-axonic synapses. (A) 2 photon z-stack reconstruction of patched L2/3 pyramidal neuron expressing either hM3Dq-mCherry or mCherry (control), filled with AlexaFluor-594 through the patch pipette (red) and the axon of a ChR2-YFP+ ChCs (green) from acute brain slices of Nkx2.1- CreERT2/ChR2+/fl mice. White arrows point at putative presynaptic boutons. (B) Representative traces of light-evoked inhibitory postsynaptic currents (IPSCs, 5x 2 ms pulses, 20 Hz) recorded from L2/3 pyramidal neurons, expressing either mCherry or the hM3Dq-mCherry, from acute brain slices of Nkx2.1-CreERT2/ChR2+/fl mice. (C) Mean amplitudes of IPSCs from L2/3 pyramidal neurons, expressing either mCherry or hM3Dq-mCherry, from acute brain slices of Nkx2.1-CreERT2/ChR2+/fl mice. Means are shown +/- SEM, statistics with non-parametric t-test with Mann-Whitney (CNO controls n = 13, hM3Dq+ n = 7, p < 0.05). (D) Mean paired-pulse ratios of IPSCs from L2/3 pyramidal neurons, expressing either mCherry or the hM3Dq-mCherry, from acute brain slices of Nkx2.1-CreERT2/ChR2+/fl mice. Means are shown +/- SEM, statistics with non-parametric t-test with Mann-Whitney (CNO controls n = 13, hM3Dq+ n = 7, p < 0.05). (E) Mean coefficient of variation (1/CV^2^) of IPSCs from L2/3 pyramidal neurons, expressing either mCherry or the hM3Dq-mCherry, from acute brain slices of Nkx2.1- CreERT2/ChR2+/fl mice. Means are shown +/- SEM, statistics with non-parametric t-test with Mann-Whitney (CNO controls n = 13, hM3Dq+ n = 7, p-value not significant). (F) Diagram explaining hypothesis on the effects of chronic increase in network activity on the nanoscale organization and function of axo-axonic synapses.

## Discussion

The current super-resolution imaging techniques that allow the localisation of molecules with nanometre precision have transformed the way we see the inner workings of cells (1). Here, we used intact brain tissue to describe how the molecules that are key for the spatial arrangement of receptors within synapses are organised into discrete sub-synaptic domains, that show distinct nanoscale motifs depending on the identity of their presynaptic partner. More importantly, these arrangements were sensitive to increases in network activity, causing changes in sub-synaptic domain volume that correlated with synapse function. Our findings show that sub-synaptic domains are a fundamental feature of inhibitory contacts that dictate the way information is transferred across the synapse.

### Arrangement of inhibitory synapses into sub-synaptic domains in intact brain tissue

We set out to characterise inhibitory synapses in intact circuits by focusing on the distribution of Gephyrin, an essential scaffolding protein in GABAergic synapses, responsible for anchoring GABA_A_R to the postsynaptic membrane. SSDs of gephyrin have been shown to co-localise with GABA_A_R domains when explored at the nanoscale (6, 7), and the complete loss of gephyrin directly affects GABA_A_R clustering (32–34). We used FingRs to tag endogenous gephyrin molecules with high signal-to-noise ratio and assess the nanoscale arrangement of GABAergic postsynaptic compartments. Using single molecule localization microscopy in brain slices, we were able to show that gephyrin is arranged in sub-synaptic domains. Although these domains are heterogenous in size and packing density, they show clear differences that depend on subcellular location and/or the presynaptic identity of the interneuron innervating it. We find that inhibitory SSDs along the dendritic tufts, which are mainly targeted by SST interneurons, were smaller, but more densely packed, than those found at the axon initial segment, which are innervated by ChCs. Understanding what determines the different nanoscale arrangement of synapses - whether postsynaptic location or presynaptic neuron identity – as well as assessing how these differences impact on synapse function, remains unknown. Differences in the nanoscale arrangement of excitatory synapses have been described along the length of dendrites in the hippocampus, or when comparing synapses formed *in vitro* onto postsynaptic neurons with different identity (either excitatory or inhibitory neurons) (35, 36). It is now becoming clear that sub-synaptic domains are key features when assessing synapse heterogeneity in the brain.

### In vivo manipulation of network activity causes a nanoscale change in inhibitory SSD size

Our most striking finding is that inhibitory sub-synaptic domains are plastic *in vivo*. Using a chemogenetic approach to increase the activity of pyramidal neurons in the cortex for two days, we found that SSDs along the AIS shrunk in volume. We saw no change in protein density per SSD, suggesting that hyperactivity caused the removal of gephyrin molecules without any change in their lattice arrangement. In contrast, we saw no change in SSD properties along dendrites, suggesting the plasticity described here was input-specific, which is in agreement with previous work showing axo-dendritic synapses formed by SST+ neurons are insensitive to chronic changes in postsynaptic activity (37). Axo-axonic synapses, on the other hand, undergo robust changes in connectivity in response to network-wide changes in activity. Previous work in the lab has shown that increases in network activity through chemogenetic stimulation during the period of axo-axonic synapse formation (P12 to P18) resulted in a marked loss of ChCs synapses at the AIS (28). Here, we implemented a similar approach for increasing cortical activity, but this time focusing on the period after synapse formation, from P20 to P22. This manipulation did not cause a change in axo-axonic synapse number. It did, however, show a reduction in SSD size, which suggests a weakened connectivity between ChCs and pyramidal neurons. Previous work in vitro has shown synapse can undergo different forms of plasticity at the nanoscale, including changes in the number of proteins per SSD (8, 21), the size of SSDs (7, 8, 21, 38, 39), the number of transsynaptic nanocolumns (7, 21), the precise alignment of transsynaptic nanocolumns (4) and the local dynamics of these proteins (21, 38, 40). Our findings provide a nanoscale description of postsynaptic inhibitory synapse plasticity *in vivo*, but many other nanoscale arrangements at the synapse still remain to be explored in intact tissue.

### Nanoscale plasticity of SSDs has a functional impact on synapse strength

A key question in the field is whether structural changes observed at the nanoscale have any functional consequence. At excitatory synapses in dissociated neurons, disruption of the nanoscale arrangement of synapses in mutant mice for genes linked to neurodevelopmental disorders, resulted in a weakening of both evoked and spontaneous synaptic transmission (41). A recent study looking at inhibitory synapses on the dendrites of neurons grown *in vitro* suggested a more nuanced view of the nanoscale arrangement of synapses. They proposed a centre-surround arrangement of receptors within the SSD that support different release modes: whereas peripheral receptors sense spontaneous release, those in the centre respond to evoked release (42). Addition of drugs that decreased the size of GABA_A_Rs SSDs through removal of receptors from the periphery of SSDs, caused a selective decrease in the amplitude of spontaneous, not evoked, events. Although we saw a decrease in the size of SSDs at the AIS, which we attribute to a loss of peripheral gephyrin, we found that this resulted in a decrease in the amplitude of evoked responses, contrary to *in vitro* findings (42). Since the plasticity we describe here was input-specific, we could not measure spontaneous inhibitory events, which would arise from all inputs onto the neuron, not just those along the AIS. Future work should explore the distinction between different modes of synaptic transmission. However, it is important to note that the functional consequences of changes in SSD properties may differ depending on preparation (e.g. dissociated neurons versus acute slices) or synapse type (e.g. axo-dendritic versus axo-axonic).

Our findings show that inhibitory synapses in the brain are indeed arranged in a plethora of different nanoscale assemblies. More importantly, these nanoscale arrangements show input-specificity, are susceptible to modulation by activity and, critically, play a central role in controlling the strength of synaptic transmission.

## Acknowledgments

We would like to thank all the members of Burrone lab for useful feedback and comments on the manuscript. We would also like to thank Nicolas Bourg and Abbelight for the SAFeRedSTORM module and software, as well as Olympus for the IX3 microscope used for dSTORM. For the purpose of open access, the authors have applied a CC BY public copyright license to any Author-Accepted Manuscript version arising from this submission.

## Author contributions

Conceptualization: BC, JB

Methodology: BC, VM, CL

Investigation: BC, VM, CL

Visualization: BC, VM, CL

Funding acquisition: BC, JB

Writing – original draft: BC, VM, CL, JB

Writing – review & editing: BC, VM, CL, JB

## Data and materials availability

All data reported in this paper will be shared by the lead contact upon request.

All original code has been deposited at https://github.com/jburrone and is publicly available as of the date of publication. Any additional information required to reanalyse the data reported in this paper is available from the lead contact upon request. Further information and requests for resources and reagents should be directed to and will be fulfilled by the lead contact, Juan Burrone.

## Supplementary Materials

**Fig. S1.**
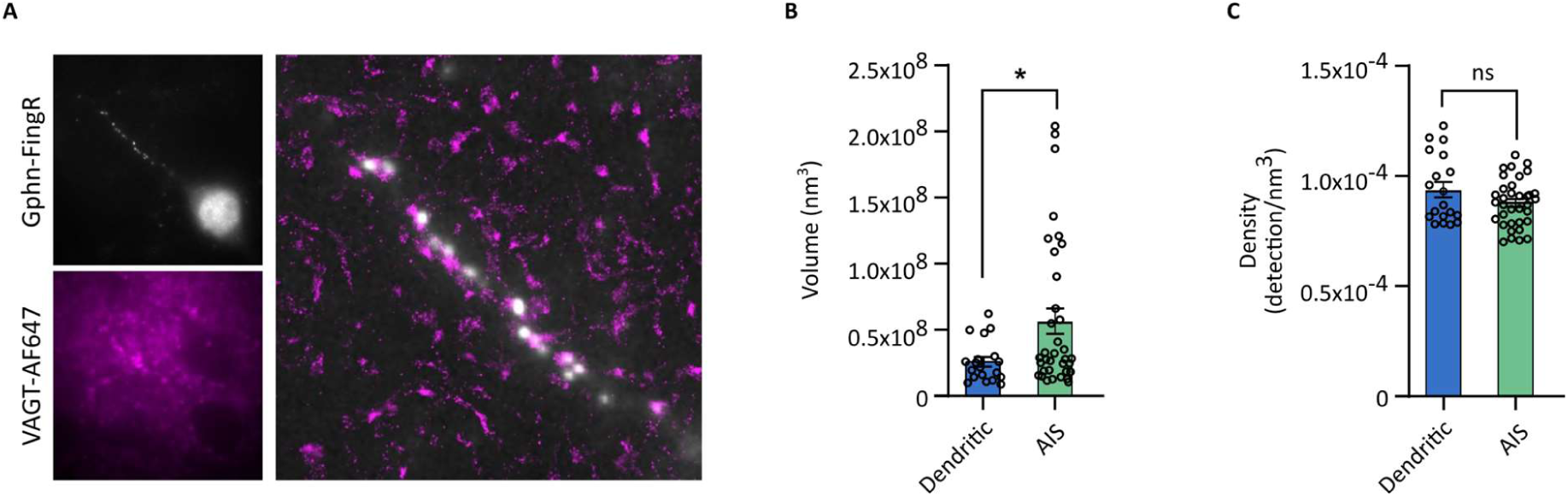
VGAT Sub-Synaptic Domains differ between axo-axonic and axo-dendritic synapses. **(A)** Widefield and 3D-dSTORM images of gephyrin-FingR and VGAT SSDs (left) and 3D-dSTORM images of VGAT SSDs (right) in axo-axonic synapses from L2/3 pyramidal neurons. Note the overlay with Gephyrin-FingR widefield. **(B)** Comparison of VGAT SSDs volume between axo-axonic and axo-dendritic synapses. Means are shown +/- SEM, statistics with non-parametric t-test with Mann-Whitney (axo-dendritic n = 35, axo-axonic n = 19, p < 0.0244). **(C)** Comparison of VGAT density per SSD between axo-axonic and axo-dendritic synapses. Means are shown +/- SEM, statistics with non-parametric t-test with Mann-Whitney (axo-dendritic n = 35, axo-axonic n = 19, p-value not significant).

**Fig S2.**
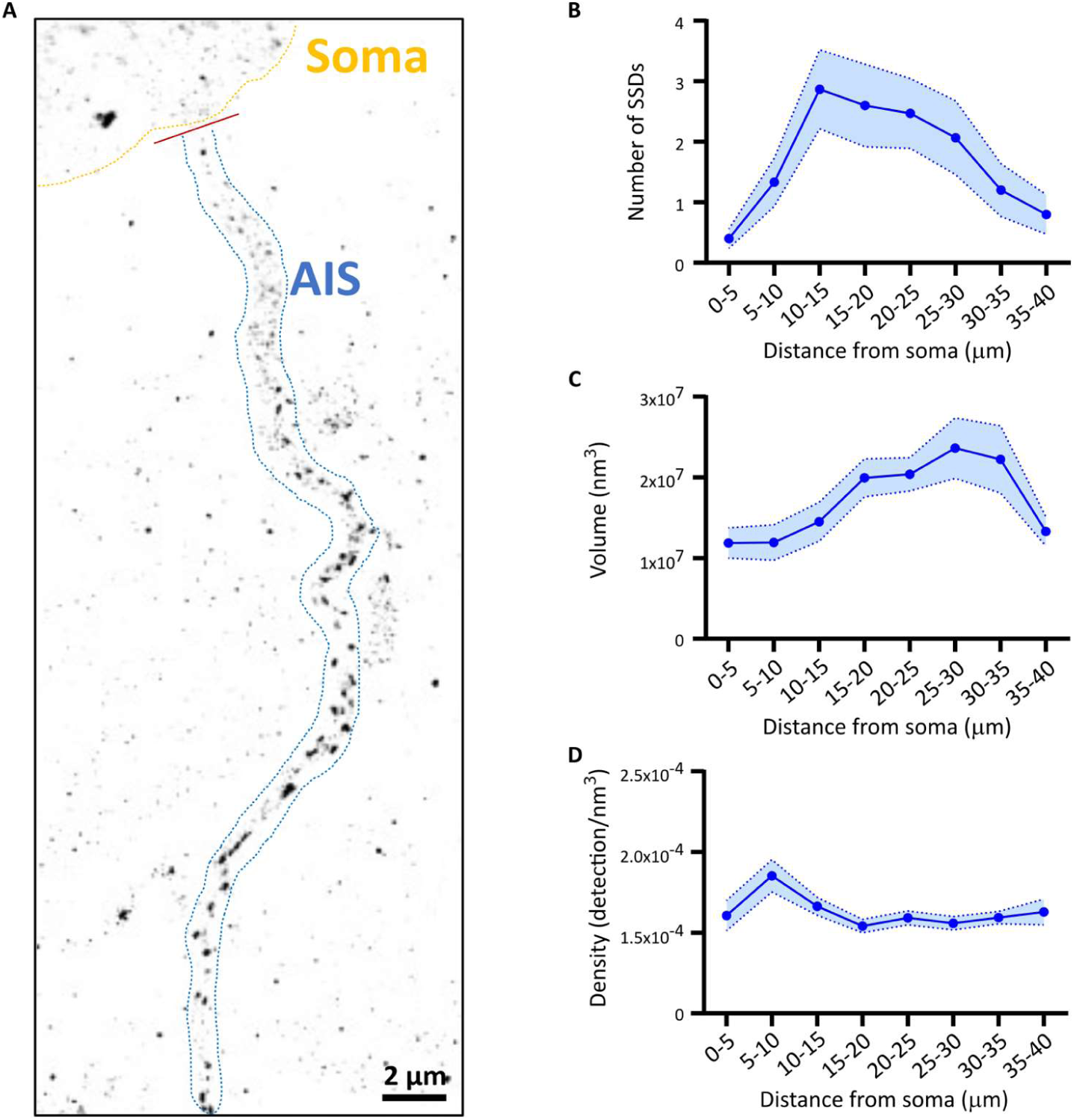
Heterogeneity of gephyrin Sub-Synaptic Domains along the AIS. **(A)** 3D-dSTORM image of gephyrin-FingR SSDs along the AIS. **(B)** Distribution of number of gephyrin clusters by the distance from the soma, at 5 μm bins. Means are shown +/- SEM. **(C)** Distribution of Gephyrin-FingR SSDs volumes along the AIS from the soma. Means are shown +/- SEM. **(D)** Distribution of Gephyrin-FingR density per SSD along the AIS, from the soma. Means are shown +/- SEM

**Fig. S3.**
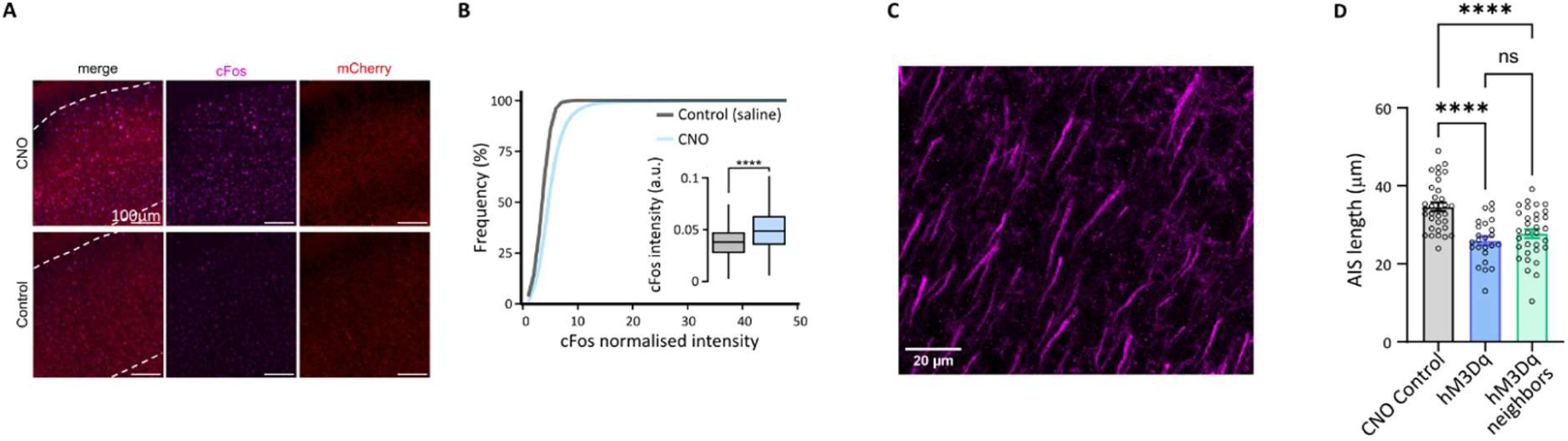
Increased neuronal activity using a chemogenetic approach. **(A)** Confocal microscopy images of cfos expression in layer 2/3 of S1 cortex, in hM3Dq-positive and hM3Dq- negative pyramidal neurons, treated with either saline or CNO. **(B)** Cumulative frequency distribution of cfos intensities from the cells in B. Inset shows mean intensities between CNO and saline groups. Means are shown +/- SEM, statistics with Wilcoxon rank-sum test (p < 0.0001, CNO n = 5491(3), Saline n = 7864(4)). **(C)** Confocal microscopy images of Ankyrin-G staining, showing the AIS labelling in layer 2/3 of S1 cortex. **(D)** Quantification of AIS lengths from neurons expressing tdTomato+ controls, hM3Dq+ and hM3Dq- neighbouring neurons. Means are shown +/- SEM, statistics with One-way ANOVA test (CNO controls n = 33, hM3Dq+ n = 24, hM3Dq- neighbours n = 32, p < 0.0001).

**Fig. S4.**
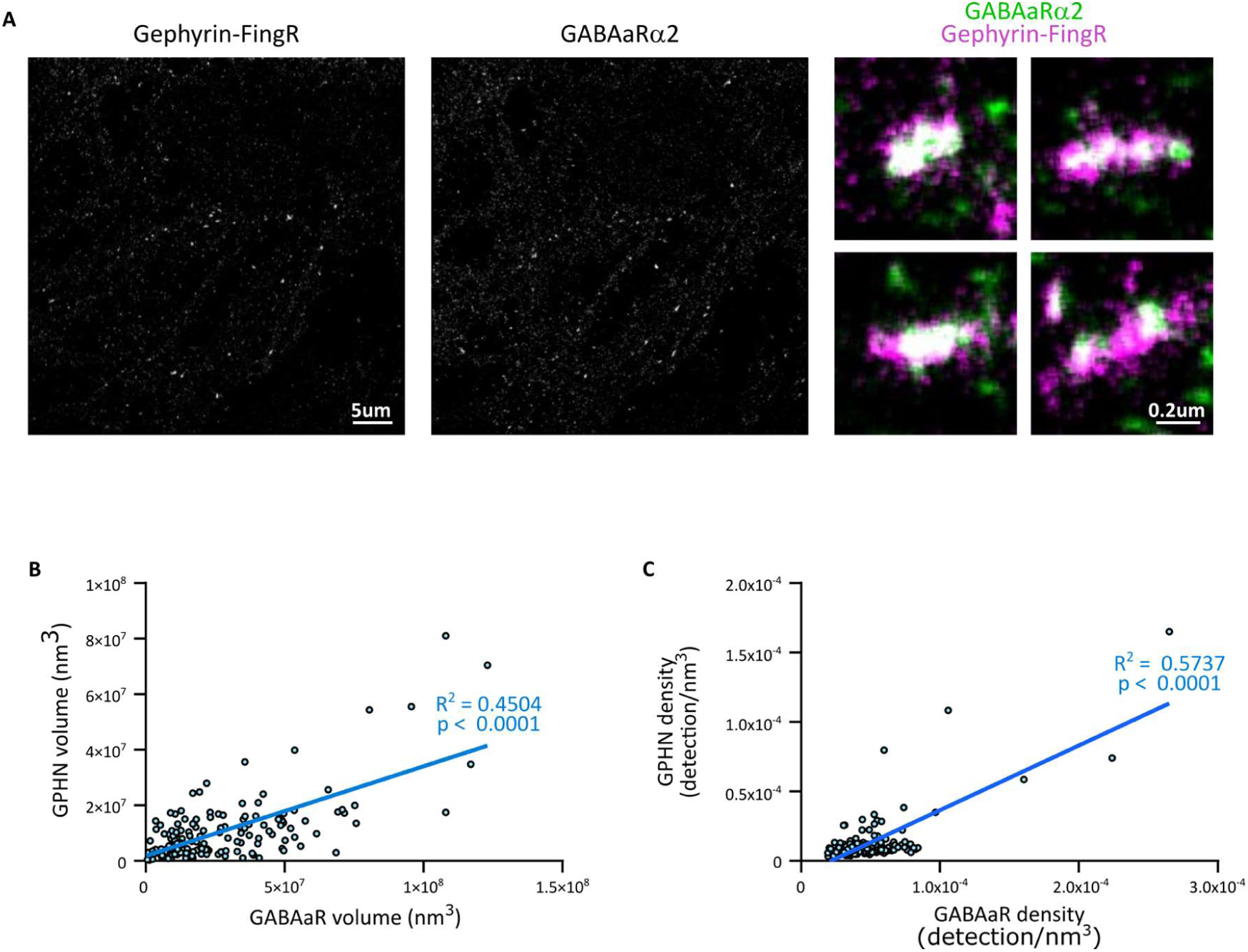
Gephyrin-FingR SSDs colocalize and co-vary with GABA_A_R-α2 SSDs. **(A)** 3D-dSTORM images of gephyrin-FingR (left) and GABA_A_R (centre) SSDs from using antibodies against GFP and GABA_A_R, respectively, in hippocampal cell cultures. Merged examples of synapses is shown on the right panel, with colocalization of GABA_A_R alpha2 (green) and Gephyrin-FingR-GFP (magenta) SSDs. **(B)** Correlative analysis GABAaR alpha2 and Gephyrin-FingR-GFP SSD volumes. **(C)** Correlative analysis of GABA_A_R alpha2 and Gephyrin-FingR-GFP SSD densities.

**Fig. S5.**
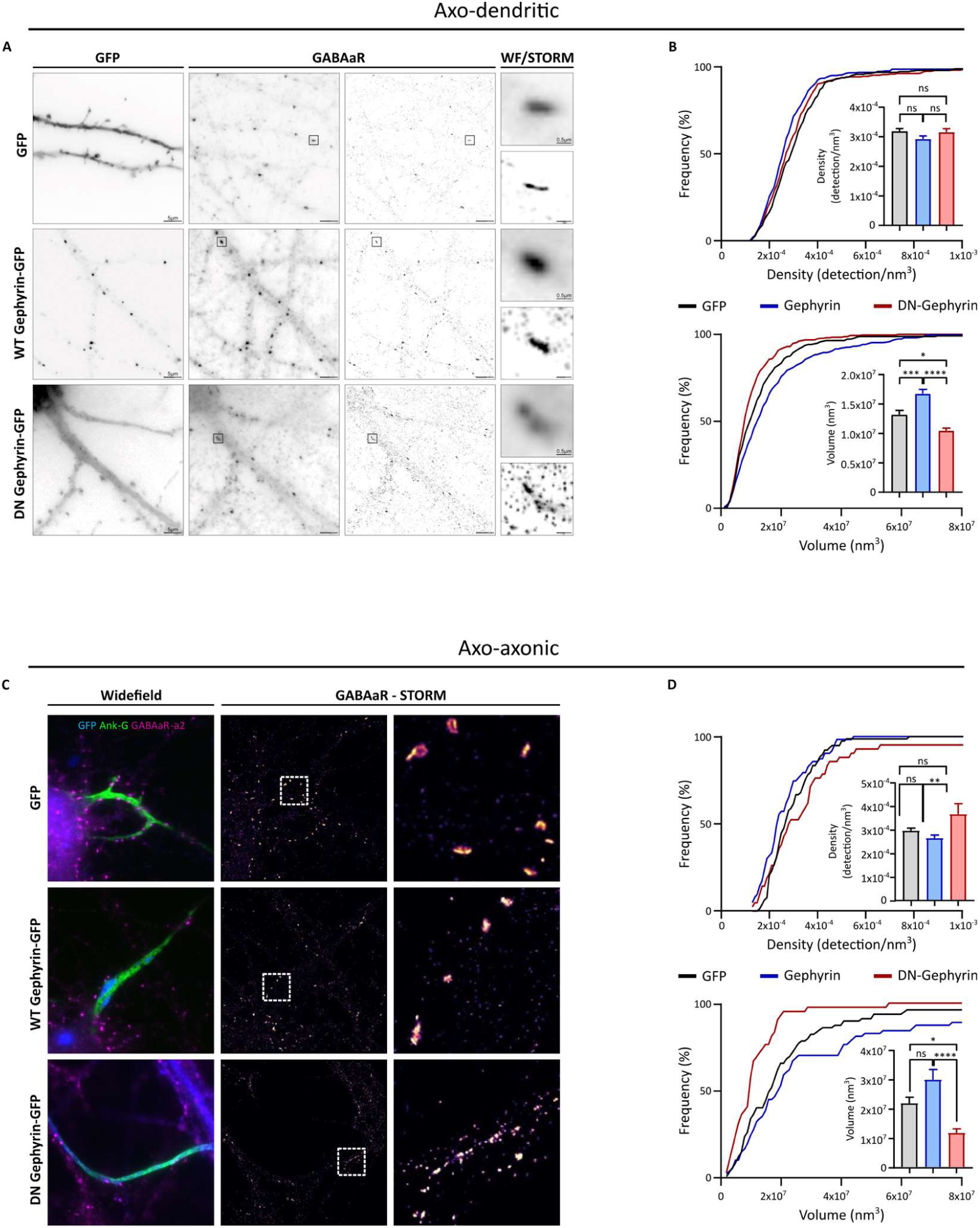
Overexpression of WT or dominant-negative Gephyrin affect GABAaR Sub-Synaptic Domains organization. **(A)** – dSTORM images of GABAaR SSD in axo-dendritic synapses from hippocampal cultured cells expressing either a WT form of gephyrin (WT-gephyrin-GFP) or a dominant-negative gephyrin variant (DN gephyrin-GFP). **(B)** - Cumulative distribution and means +/- SEM (inset) of axo-dendritic synapses from the conditions in A showing GABAaR densities per SSD (top, statistical analysis with one-way ANOVA with Tukey’s multiple comparisons test (GFP (n = 259) vs DN-gephyrin-GFP (n = 315), p-value non-significant; GFP (n = 259) vs WT-gephyrin-GFP (n = 384), p-value non-significant; WT-gephyrin-GFP (n = 384) vs DN-gephyrin-GFP (n = 315), p-value non-significant) and SSDs volumes (bottom, statistical analysis with one-way ANOVA with Tukey’s multiple comparisons test (GFP (n = 259) vs DN-gephyrin-GFP (n = 315), p < 0.05; GFP (n = 259) vs WT-gephyrin-GFP (n = 384), p < 0.001; WT-gephyrin-GFP (n = 384) vs DN-gephyrin-GFP (n = 315),p < 0.0001). **(C)** dSTORM images of GABAaR SSD in axo-axonic synapses from hippocampal cultured cells expressing either a WT form of gephyrin (WT-gephyrin-GFP) or a dominant-negative gephyrin variant (DN gephyrin-GFP). **(D)** Cumulative distribution and means +/- SEM (inset) of axo-axonic synapses from the conditions in A showing GABAaR densities per SSD (top, statistical analysis with one-way ANOVA with Tukey’s multiple comparisons test (GFP (n = 78) vs DN-gephyrin-GFP (n = 42), p-value non-significant; GFP (n = 78) vs WT-gephyrin-GFP (n = 63), p-value non-significant; WT-gephyrin-GFP (n = 63) vs DN-gephyrin-GFP (n = 42), p < 0.01) and SSDs volumes (bottom, statistical analysis with one-way ANOVA with Tukey’s multiple comparisons test (GFP (n = 78) vs DN-gephyrin-GFP (n = 42), p < 0.05; GFP (n = 78) vs WT-gephyrin-GFP (n = 63), p-value non-significant; WT-gephyrin-GFP (n = 63) vs DN-gephyrin-GFP (n = 42), p < 0.0001).

**Fig. S6.**
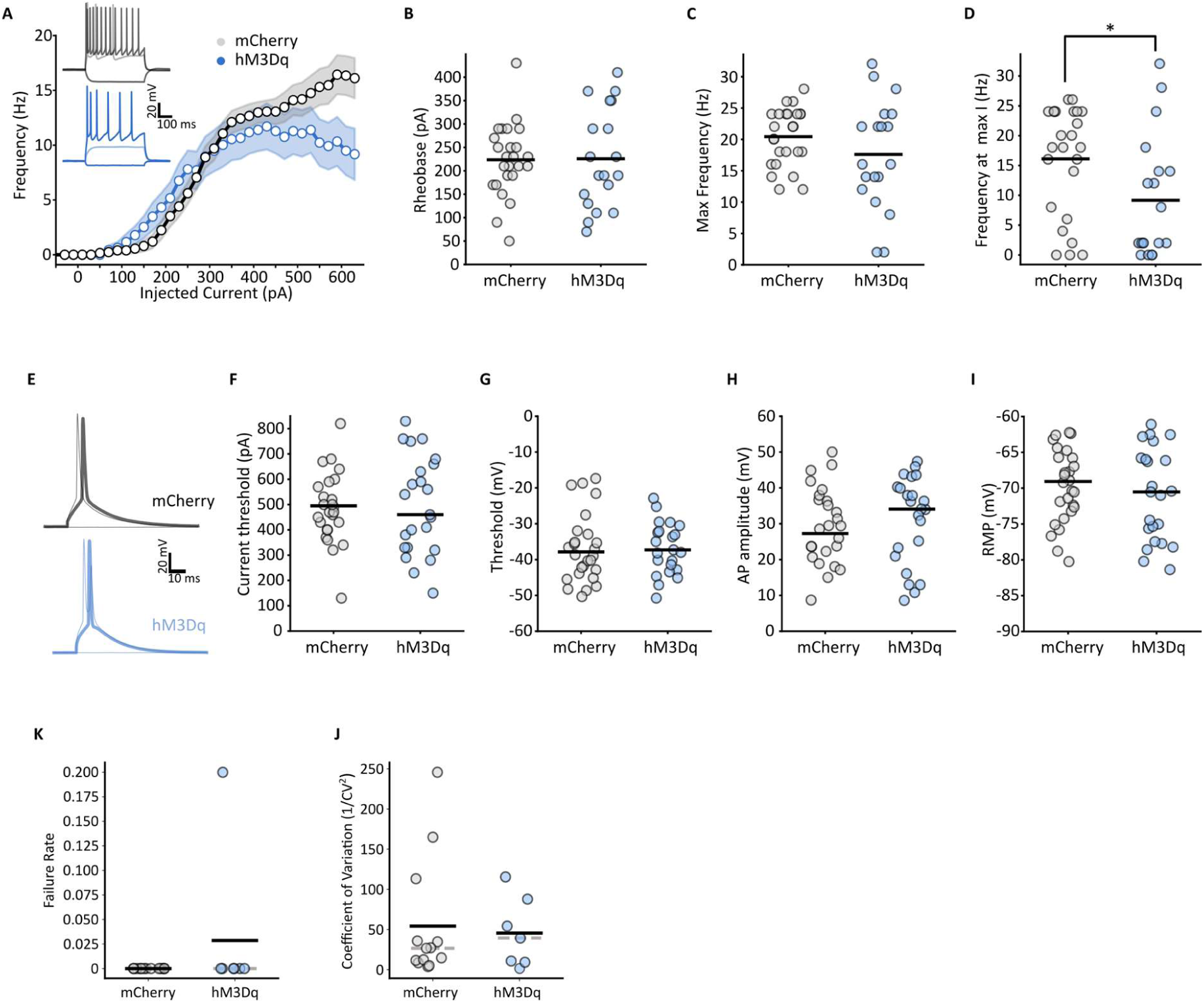
Intrinsic properties of hM3Dq-activated pyramidal neurons. **(A)** Input-output plot of action potentials firing frequency from current clamp recordings (−150 to 400 pA, 20 pA increments, 500 ms) of hM3Dq-positive or mCherry-positive (controls) L2/3 pyramidal neurons from acute brain slices of mice treated with CNO. Insets show representative voltage traces at the same current injection steps (0, 100, 600 pA). **(B-D)** Rheobase, Max frequency and frequency at the last current step from the voltage recordings in A. **(E)** Representative voltage traces for current clamp recordings (0 to 900 pA, 10 pA increments, 10 ms) of single action potentials of hM3Dq-positive or mCherry-positive (controls) L2/3 pyramidal neurons from acute brain slices of mice treated with CNO. **(F-I)** Current threshold of firing, voltage threshold, action potential amplitude and resting membrane potentials from the voltage recordings in E. **(K-J)** Failure rate and coefficient of variation from the eIPSCs recordings (5x 2 ms pulses, 20 Hz) recorded from L2/3 pyramidal neurons, expressing either mCherry or the hM3Dq-mCherry, from acute brain slices of Nkx2.1-CreERT2/ChR2+/fl mice.

## Materials and Methods

### For in vivo animal studies

All animal procedures were approved by the local ethics committee and licensed under the UK Animals (Scientific Procedures) Act of 1986. Male and Female CD1, C57BL/6-ChR2-floxed and Nkx2-1CreERT2 mice were housed grouped in standard cages and provided with ad libitum food and water. CD1 wildtype mice were used for all STORM animal experiments, aged between P20 and P22 (as indicated in the manuscript). A cross between C57BL/6-ChR2-floxed and Nkx2-1CreERT2 mouse lines aged between P20 and P22 (as indicated in the manuscript) was used for all electrophysiology experiments. For chemogenetic experiments, mice were treated with Clozapine-N-Oxide (CNO, 1 mg/kg) twice a day for 2 consecutive days.

### Neonatal intracranial virus injection

Mouse pups age P1 were anaesthetized with isoflurane (3%) and immobilized on a stereotaxic frame (World Precision Instruments) adapted for mouse pups. Borosilicate capillary needles were filled with a mixture of adeno-associated viruses (AAVs). AAVs for the expression of Gephyrin-FingR (AAV8-EF1A-FLExFRT-Gephyrin.FingR-eGFP-CCR5TC) and hM3Dq (pAAV-CaMKIIa- hM3D(Gq)-mCherry) were used for confocal and STORM microscopy experiments, while only the latter was used for electrophysiological experiments. Injection coordinates to target L2/3 of the somatosensory cortex were A/P 1.6, M/L 1.5, D/V 0.1 from lambda.

### Dissociated neuron preparation

Primary hippocampal cultured neurons were prepared from Wistar rat embryos at E18. Dams were culled via a schedule 1 procedure according to the Animals (Scientific Procedures) Act 1986. Sex of cells was not recorded, due to the developmental timepoint of the embryos. Hippocampi were dissected, and incubated with trypsin (Worthington’s, 0.33 mg/ml) for 15-min at 37°C. Three fire-polished pipettes of diminishing diameters were used for mechanical dissociation. Neurons were pelleted at 200RCF for 10-min, then washed in Hanks’ Balanced Salt Solution (Life Technologies). Neurons were suspended in neurobasal media with 2% B27 (Life Technologies), 2% fetal bovine serum (FBS), 1% glutamax and 1% penicillin streptomycin. Neurons were plated onto glass cover slips (Menzel Gläser, Germany) at a density of 90,000 neurons per coverslip.

### Transfection and culture fixation

Primary hippocampal cultured neurons were transfected using Lipofectamine 2000 transfection reagent (Thermofisher) diluted in complete neurobasal media with the plasmid DNA (1.25μg per 25μL media/coverslip) for 25-min at 37°C. Neurons were fixed by incubating with 4% PFA (in pbs) for 5-min at room temperature.

### Acute Slice Preparation

Nkx-2.1-CreERT2/ChR2-floxed mice (age P22) were anaesthetised with Ketamine (200mg/kg) and Xylazine (40mg/kg) and decapitated. Brains were quickly harvested and kept in cold slicing solution (93 mM NMDG, 20 mM Glucose, 20 mM Hepes, 2.5 mM KCl, 30 mM NaHCO2, 1.2 mM NaH2PO4, 7 mM Na-Ascorbate, 2 mM Thiourea, 4 mM Na-Pyruvate, 0.5 mM CaCl2, 10 mM MgCl2, pH 7.35 carbonated with 95% O2/5% CO2) for 1-2 minutes. Brains were transferred on a vibratome (7000smz-2, Campden Instruments, UK) and coronal mouse brain slices (300 mm thickness, speed 0.05 mm/s, 70 Hz/1 mm amplitudes) were collected and incubated for 20 minutes in carbonated slicing solution at 32° C. Slices were then transferred in holding solution (95 mM NaCl, 12 mM Glucose, 20 mM Hepes, 2.5 mM KCl, 30 mM NaHCO2, 1.2 mM NaH2PO4, 7 mM Na-Ascorbate, 2 mM Thiourea, 4 mM Na-Pyruvate, 2 mM CaCl2, 2 mM MgCl2, pH 7.35 carbonated with 95% O2/5% CO2) at room temperature and let recover for at least 2 hours prior to experiments.

### Electrophysiology

After recovery, coronal mouse brain acute slices from Nkx-2.1-CreERT2/ChR2-floxed mice were transferred to a recording chamber equipped with a custom-built microscope (Cosys Ltd) supplied with Dodt gradient contrast and infrared light and superfused with recording aCSF solution (124 mM NaCl, 5 mM KCl, 1.25 mM NaH2PO4, 6 mM Glucose, 26 mM NaHCO3, 5 mM HEPES, 2 mM CaCl2, 1 mM MgCl2, pH 7.35 carbonated with 95% O2/5% CO2 at 32°C. Traces were acquired with a Multiclamp 700B amplifier (Axon, Molecular Devices) and digitised with the Digidata 1550B digitizer (Axon, Molecular Devices) with 10 kHz sampling rate and 5 kHz low-pass filter. Data was acquired with the software Clampex 10.7 (Axon, Molecular Devices).

For light-evoked inhibitory postsynaptic currents (IPSCs), primary somatosensory cortex L2/3 pyramidal neurons expressing hM3Dq-mCherry and in proximity (200 mm radius) to ChR2-YFP- positive chandelier cells (ChCs) were selected. Whole-cell configuration was achieved using a borosilicate pipette (World Precision Instruments) with a resistance of 4-5 MOhm and containing internal solution (120 mM CsCl, 5 mM KCl, 10 mM HEPES, 0.5 mM EGTA, 5 mM MgATP, 0.3 mM NaGTP, 5 mM Na2-Phosphocreatinine, 1 mM MgCl2, 5 mM QX-314, 20 μM AlexaFluor 594, pH 7.3), Light-evoked IPSCs were recorded from the patched pyramidal neuron in voltage clamp while holding the neuron at –70 mV. Recording aCSF was supplemented with the addition of 50 mM APV and 2 mM CNQX, to eliminate. ChR2-YFP+ ChCs with the potential of forming synapses onto the patched neuron were stimulated using 5 light pulses at 20 Hz, 2 ms long, with a 470 nm fluorescent light at 4 mW power. All data was analysed and plotted using a custom-made Python script.

### Histology for dSTORM and confocal

Mice (CD1, P22) were anaesthetized with an overdose of sodium pentobarbital and transcardially perfused with 40 mL of ice-cold saline solution followed by 40 mL of 4% (w/v) PFA (pH 7.3). The brains were carefully removed and put in 4% (w/v) PFA solution overnight at 4°C. Brains were incubated in 15% sucrose then in 30% sucrose, both overnight at 4°C. They were embedded in blocks of Gelatine/Sucrose and froze in isopentane (−60°C) and then stored at −80°C until cryo-sectioning. Coronal slice of 40µm thick were obtained with a cryostat (Leica). Free-floating slices were kept in anti-freeze PBS at −20°C. On the day of staining, slices were incubated with warm PBS 4 times 5 minutes each, to remove the gelatine/sucrose blocks. A 50mM NH4Cl solution was used to quench the remaining PFA. For permeabilization, sections were incubated 4 times in a 0.25% Triton X-100 solution for 15 minutes each at room temperature (RT) on a shaker. Brain slices were incubated in a Blocking Buffer (5% BSA, 0.25% Triton X100, 10% Goat Serum) for 2 hours at RT on a shaker. This was followed by an overnight incubation with primary antibodies diluted in an Antibody Buffer (1% BSA, 0.25% Triston X-100, 5% Goat serum). On the following morning, slices underwent four 15 minutes PBS-Triton X-100 (0.25%) washes, followed by a 2- hour incubation at room temperature with secondary antibodies diluted in the Antibody Buffer. The secondary antibody solution was washed off as before and slices were mounted onto 1.5H 25mm glass coverslip (Marienfeld). Slices were allowed to dry on the coverslips, melted 2% agarose was used to immobilize the tissue, and coverslip were kept in PBS at 4°C until imaging.

### 3D dSTORM imaging

For dSTORM, coverslips were placed in an imaging chamber (AttoFluor Cell Chamber, Invitrogen). The imaging chamber was filled with STORM Buffer and sealed using another glass coverslip. Final dSTORM imaging buffers were composed of oxygen scavengers (100 μg/ml Glucose oxidase (Sigma-Aldrich G2133) and 4 μg/ml Catalase (Sigma-Aldrich C100)) and reductive agent (100mM β-Mercaptoethylamine-HCl, Sigma-Aldrich M6500). Oxygen scavengers stock solution containing 20mM Tris–HCl pH 7.2, 4mM TCEP, 25mM KCl, 50% Glycerol and 1mg/ml Glucose oxidase (Sigma-Aldrich G2133) and 42 μg/ml Catalase (Sigma-Aldrich C100) and was stored at −20°C. Reducing agent stock solution containing 1M β-Mercaptoethylamine-HCl (Sigma-Aldrich M6500) in deionized water and pH adjusted at pH 8 with NaOH, was stored at −20°C. Dilution STORM Buffer containing 100mg/mL Glucose (Sigma-Aldrich) and 10% Glycerol (Fisher Scientific) in deionized water, was stored at 4°C.

Single color dSTORM imaging was performed using a Nikon N-STORM5 (Nikon, Japan) equipped with a silicon oil-immersion objective (100X 1.35NA silicon oil immersion, Nikon). To achieve 3D, a cylindrical lens was inserted in front of the ORCA-Flash4.0 sCMOS camera (Hamamatsu). Image acquisition and control of microscope were driven by Nikon NIS element software.

Multicolor dSTORM imaging was performed using a spectral-demixing SAFeRedSTORM module (Abbelight, France) mounted on an Evident/Olympus IX3 equipped with an oil-immersion objective (100× 1.5NA oil immersion, Evident/Olympus) and fiber-coupled 642nm laser (450mW Errol, France). Fluorescent signal was collected with two ORCA-Fusion sCMOS camera (Hamamatsu). A long-pass dichroic beam splitter (700nm; Chroma Technology) was used to split the emission light on the two cameras. 3D-dSTORM imaging was performed by using cylindrical lenses placed before each camera.

Image acquisition and control of microscope were driven by Abbelight’s NEO software. The image stack contained 60,000 frames. Selected ROI (region of interest) had dimension of 512 × 512 pixels (pixel size = 97 nm). Cross-correlation was used to correct for lateral drifts and chromatic aberration was avoided using spectral demixing technique. Super-resolution images (.tiff) with a pixel size of 10 nm and localization files (.csv) were obtained using NEO_analysis software (Abbelight).

### 3D-dSTORM analysis

Point Clouds Analyst software (PoCA, https://poca-smlm.github.io/) (39) was used to identify and analyze Sub-Synaptic Domains from localized molecule coordinates (.csv) of FingR-Gephyrin, VGAT or GABAaR, as described in Levet et al 2015(43). First a Voronoi diagram was applied to draw 3D polygons of various sizes centered on the localized molecules. Second, an automatic segmentation was performed to detect a “single fluorophore” (which can be an isolated antibody or isolated protein) and estimate the number of detections per fluorophore. Then, another automatic segmentation was performed to detect clusters for each protein of interest. SSDs were thresholded based on their density (δi1 for single color dSTORM experiment, or δi1 and δi2 for color1 and color2 respectively in the case of dual color dSTORM experiments) with respect to the average density of the image for each color (δd1 or δd2), such that δi1 ≥ 1δd1 and δi2 ≥ 1δd2. All selected neighboring molecules were considered as forming a cluster when having a minimum number of 10 or 20 times the median value of “single fluorophore” count for FingR-Gephyrin if labeled with nanobody (figure 3) or primary/secondary antibody (figure 1) respectively, 30 times the median value of “single fluorophore” count for VGAT and 10 times the median value of “single fluorophore” count for GABAaR alpha2.

### Confocal microscopy

Mounted slices were imaged via a Nikon Ti-E inverted equipped with an A1R confocal microscope. Neurons were imaged with a 60x oil objectives using the 640nm, 561nm, 488nm and 405nm laser lines and collected with the appropriate emission filters. Z-stacks were acquired, with 0.175µm steps. Typical pixel size (X/Y) was 0.11µm.

### Analysis of confocal images

Axons were traced using Simple Neurite Tracer (ImageJ Plugin66) based on mCherry or tdTomato staining (from hm3dq-mcherry or soluble tdTomato) and exported as swc files. Axon Initial Segment and FingR-Gephyrin-GFP puncta were automatically detected using a trained model (Weka plug-in Fiji). Analysis of the AIS length, number of FingR-Gephyrin and number of VGAT puncta along each AIS was undertaken in IgorPro (version 6.37) using custom-built scripts (courtesy of Dr Guilherme Neves). Maximum Fluorescence intensity and the centroid of each ROI were measured in all 3D gaussian filtered (radius = 2 pixels) confocal slices. ROI coordinates were used to determine the number of axo-axonic synapses within the swc trace.

### Quantification and statistical analysis

All statistical details of experiments can be found in Table 1 and in the figure legends. All electrophysiological data was analysed and plotted using a custom-made Python script. Graphpad Prism (version 9.1.1) was used for statistical analysis of the confocal and dSTORM. Correlations were assessed using Spearman’s rank correlation. Comparisons between group pairs were tested using non-parametric t-test or Wilcoxon rank-sum test. Comparisons between multiple groups were tested using one-way ANOVA tests.Type or paste text here. You can break this section up into subheads as needed (e.g., one on “Materials” and one on “Methods”).

**Table S1.** Reagents and resources.

| Reagent/Resource | Source | Identifier |
| --- | --- | --- |
| **Antibodies** |  |  |
| Anti-GFP FluoTag-X4 nanobody coupled with sulfoCy5 (Dil. 1/500) | Nanotag |  |
| Anti-GFP FluoTag-X4 nanobody coupled to AlexaFluor647 (Dil. 1/500) | Nanotag |  |
| Anti-GFP FluoTag-X4 nanobody coupled to Atto488 | Nanotag |  |
| Anti-RFP FluoTag-X4 nanobody coupled to AZDye568 (Dil. 1/500) | Nanotag |  |
| Anti-GFP antibody (Dil. 1/2000) | Abcam | ab13970 |
| Anti-VGAT (Rabbit, Dil. 1/1000) | Synaptic System | 131 002 |
| Anti-VGAT (Mouse, Dil. 1/500) | Synaptic System | 131 011 |
| Anti-GABAαR alpha (Rabbit, Dil. 1/1000) | Synaptic System | 224 103 |
| Anti-AnkyrinG (Mouse, Dil. 1/500) | NeuroMab | 75-146 |
| Anti-cfos (Rabbit, Dil. 1/1000) | Abcam | Ab190286 |
| Anti-mouse AlexaFluor647+ (Goat, Dil. 1/1000) | Invitrogen | A32728 |
| Anti-rabbit AlexaFluor647 (Goat, Dil. 1/1000) | Invitrogen | A21245 |
| Anti-rabbit CF680 (Goat, Dil. 1/1000) | Biotium | 20818 |
| **Viruses** |  |  |
| AAV8-EF1A-FLEXFRT-Gephyrin.FingR-eGFP-CCR5TC | Horton et al. 2024 |  |
| pAAV-CaMKIIα-hM3D(Gq)-mCherry | Addgene | 50476-AAV9 |
| **Software** |  |  |
| GraphPad Prism | Graphpad | https://www.graphpad.com/features |
| Point Clouds Analyst software (PoCA) |  | https://poca-smlm.github.io/ |
| Python | Python.org | https://www.python.org/ |

